# Effects of bacterial wilt on community composition and diversity of culturable endophytic fungi in *Alpinia galanga*

**DOI:** 10.1101/2024.11.23.625029

**Authors:** Liu Jiahui, Wu Yuanyuan, Lin Jinru, Xie Mengxia, Chen Likai, Wang Liguo

**Affiliations:** Laboratory of Germplasm Resources and Molecular Identification of Traditional Chinese Medicine, School of Pharmaceutical Sciences, Guangzhou University of Chinese Medicine, Guangzhou, China; Artemisinin Research Center, Guangzhou University of Chinese Medicine, Guangzhou, China; Key Laboratory of Lingnan Chinese Medicine Resources, Ministry of Education, Guangzhou University of Chinese Medicine, Guangzhou 510006, China

## Abstract

Hongdoukou plant (*Alpinia galanga* Willd.) is a perennial herbaceous plant that usually has a stable microflora living in the inter-root and stem and leaf tissues, which assists the host in normal growth and development. The bacterial wilt disease that had been investigated in planting bases of *A. galanga*, was a new soil-borne disease infected by pathogenic bacteria (*Ralstonia solanacearum* (Smith) Yabuuchi et al.), which disrupts the *A. galanga*-microbe-soil microecological balance. For this reason, it is important to study the changes of the community composition and diversity of endophytic fungi in healthy and diseased *A. galanga* (HDK_J and HDK_B), and to mine the active endophytic fungal resources in order to lay the foundation for exploring the functional microbial communities for artificial synthesis. All experimental materials were from the healthy and diseased plant clusters of *A. galanga* in the traditional planting areas in Guangdong Province. The endophytic fungi were isolated from the stems and leaves of *A. galanga* by separated plant tissue blocks cultured in PDA medium. The colonization rate (CR) was calculated, and the isolated strains were amplified by PCR with the ITS1/ITS4 genes as the fungal molecular markers, to identify the endophytic fungi species, and to analyze the diversity of endophytic fungi in the healthy and diseased clusters of HDK_J and HDK_B, and the indexes included: relative segregation frequency (SF), relative abundance (RA) plotted by ggplot2 under R language, Shannon-Wiener diversity index (*H′*), Simpson index (*D*), Margalef richness index (*R*) and Pielou uniformity index (*J*), and finally, a phylogenetic tree of the endophytic fungi of healthy and diseased *A. galanga* was constructed by MEGA. A total of 685 endophytic fungal strains were isolated from the stems and leaves of healthy (HDK_J) and diseased (HDK_B) *A. galanga*, of which 27 endophytic fungal species were identified in healthy *A. galanga*. and 8 in diseased *A. galanga*, which belonged to 3 Phyla, 6 Classes and 13 Families. At the genus level, the RA of HDK_B endophytic fungi were only 38.93% of HDK_J, with significant difference (*P*<0.05), indicating that the composition and abundance of endophytic fungi in HDK_B were lower than that in HDK_J. Meanwhile, the endophytic fungi species isolated from the stems and leaves of the HDK_J and HDK_B were the same, but the RA of leaf organs was higher than that of stems, with the RA in the leaves in HDK_J higher than that of stems by 124.23%, and the RA of leaves in HDK_B higher than that of stems by 78.23%. The RA of HDK_J leaves was 78.08% higher than that of stems. all the diversity indexes of HDK_J were higher than that of HDK_B, with significant differences (*P*<0.05), and the Shannon-Wiener index (*H′*), Simpson’s index (*D*), and Margalef’s index (*R*) behaved as follows: HDK_J leaves (1.16, 0.87, 5.12) > HDK_J stems (0.82, 0.72, 4.22) > HDK_B leaves (0.57, 0.24, 2.97) > HDK_B stems (0.33, 0.14, 2.25), whereas Pielou’s index (*J*) indicated that the endophytic fungal species were more uniformly distributed in HDK_J leaves, and there was not much difference in the homogeneity of the leaves of the HDK_J stems and the HDK_B stems. The mapping phylogenetic trees showed there were 4 major development branches of the species on endophytic fungi in HDK_J, and especially, there were many long development branches under the Ascomycota. The genetic relationships among different species were completely revealed in phylogenetic trees. The average genetic similarity among genera was 52.80%. Conversely, there only were1major branch (Ascomycota), and few sub-branches under the Ascomycota in HDK_B. The average genetic similarity among genera was 84.59%, and the difference was small. The bacterial wilt significantly affected the composition and RA of endophytic fungi in *A. galanga*. The diversity indices showed decreasing trend in *A. galanga* after infected by *R. solanacearum*. The dominant species were changed. The parts of sensitive endophytic fungi were disappeared. This result will be helpful for the study on the relationship between the artificial minimal microbial community and the role of the host as well as the study on synthetic microbiomics.

## Introduction

Plant growth is susceptible to various biotic and abiotic stress factors, and the normal development process of plants is impeded under the stress, which affects the traits and quality of plants. The stressed plants not only be caused by the pests and diseases, but also be caused by abiotic factors such as drought, waterlogging, soil salinity, air pollution, excessive heavy metals and nutrient deficiency. Stressed plants trigger a series of responses in the host, including changes in various physiological and biochemical indices in metabolic pathways and adjustments in the microenvironment of host tissue system which the microbes live in [1]. There are a lot of endophytes living in plant tissues, and their community structure is closely related to the hosts and the ecological environments. When plants live in a specific geographical environment for a long time, there are often stable endophyte community systems in the plant tissues. The endophytic flora co-exist and interact with the host for a long period of time, and the small molecule metabolites produced by either the host or the endophytes during the interaction play a key role in regulating the growth and development of the host, the defence of the system, and the synthesis and accumulation of the secondary metabolites [2,3,4]. After healthy plants are infected with the diseases, the balance microenvironment of hosts was disrupted in vivo, leading to the changes of the endophyte community and the abundance of the endophyte constitution, and some inferior endophytic flora become dominant flora, and some endophytes disappear, and some endophytes become dominant endophytes which affect the functions of endophyte community [5]. An increasing number of studies on endophytes have shown the diversity of the endophytes functions which are not identical for different active endophytes, and competitive endophytes have ability to compete with pathogens for to obtain nutrients, to occupy invasion sites and to resist the invasion of pathogens, and antagonistic endophytes are able to inhibit the expansion and spread of pathogens in host tissues [6]. With the fluctuation of the endophyte community of the hosts, the normal interspecific coexistence, mutuality, symbiosis, parasitism, competition and antagonism between the endophyte community and the exogenous pathogenic microorganisms are changed, especially the changes of the endophyte community composition with competitive or antagonistic ability will affect the development of the host diseases. More and more studies have been found that the community structure of endophytes is related to the disease resistance, the pest resistance and stress resistance in hosts [7,8,9]. By studying the changes in the structural composition and diversity of endophyte communities in healthy and infected hosts, a foundation can be laid for exploring the construction of recombinant artificial endophytes and their functions.

Hongdoukou is a perennial herb in the ginger family (Zingiberaceae), and its mature dried fruits and rhizomes are two different chinese medicinal materials, which are commonly used medicinal and food herb cultivated in Guangdong, Guangxi and Yunnan Provinces in China. In recent years, bacterial wilt has become a serious disease threatening the growth of *A. galanga*. Our research team first identified the causative agent causing bacterial wilt according to Koch’s rule, expanding the host range of pathogenic bacteria. Bacterial wilt is a bacterial soilborne disease caused by *Ralstonia solanacearum*, which can cause damage in many tropical and subtropical crops. Bacterial wilt is a typical vascular disease, in which *R. solanacearum* invades the host from the root wounds, passes through the cortex into the vascular bundles, and spreads upward along the vascular bundles, during which the mass bacterial propagules and secretions adhere to the inner wall of the conduit to form a biofilm barrier to block the conduit, causing leaves to lose water and green withering [10]. Earlier, the studies of bacterial wilt mainly focused on the interacting mechanism between pathogenic process of *R. solanacearum* and host resistance, and it was detected that *R. solanacearum* secretes extracellular polysaccharides (EPS) [11] and a variety of pathogenicity-associated effector proteins (EPs) [12], which are virulence factors that determine the pathogenicity and host range of *R. solanacearum* [13,14]. Meanwhile, the signaling channels of the innate immune system of host had also been studied from the perspective of host-pathogen interaction, and quite a few of signaling components and cell membrane surface pattern-recognition receptors (PRRs) had been discovered and examined [15,16].In recent years, with the development of multi-omics analysis techniques, the rhizobial microbiome and endophytic flora that can help the host to resist bacterial wilt have been increasingly reported, and a number of useful bacteria and fungi successively have been found, and it can indicate that the plant immune system and microbiomes interact synergistically with each other to defend against *R. solanacearum* [17,18]. In our previous study on the community structure of endophytic fungi, their stability was greatly affected by the external environment, and it was hypothesized that the changes in the host tissue microenvironment would affect the endophytic fungal community composition and diversity before and after *A. galanga* is infested by *R. solanacearum*. Therefore, we have isolated endophytic fungi from the both healthy and diseased materials of *A. galanga* with traditional fungal PDA culture method to study the differences in the composition of the culturable endophytic fungal community and the changes of diversity in the two materials. Furthermore, it is aim for to explore the synergistic role of endophytic fungi and the host immune system in defence against bacterial wilt.

## Materials and Methods

### Source of Materials

From March 2022 to May 2023, diseased samples of *A. galanga* (No. HDK_B) were collected from the growing regions that is managed by an agricultural company in Chaozhou City in China (Figure 1 A). According to the epidemic characteristics of soilborne diseases, healthy *A. galanga* (No. HDK_J) were collected from distant mountainous regions far away from the pathogen-contaminated soils in Chaozhou City in China’s Guangdong Province. (Figure 1 B). Healthy and diseased samples were collected in batches at three growth stages approximately five months apart, at which time the new bacterial wilt spreads from plant to plant. 1 branch was cut from each plant cluster, and 5 branches were collected from each healthy and diseased *A. galanga*. After all samples were taken back to the laboratory, the stems (No. HDK_S) and leaves (No. HDK_L) of whole branches were separated, rinsed with sterile water and dried. Then, all stems and leaves were clipped 10 cm size, and packaged in sealing bags, and stored in refrigerator at 4 °C for later.

**Figure 1.**
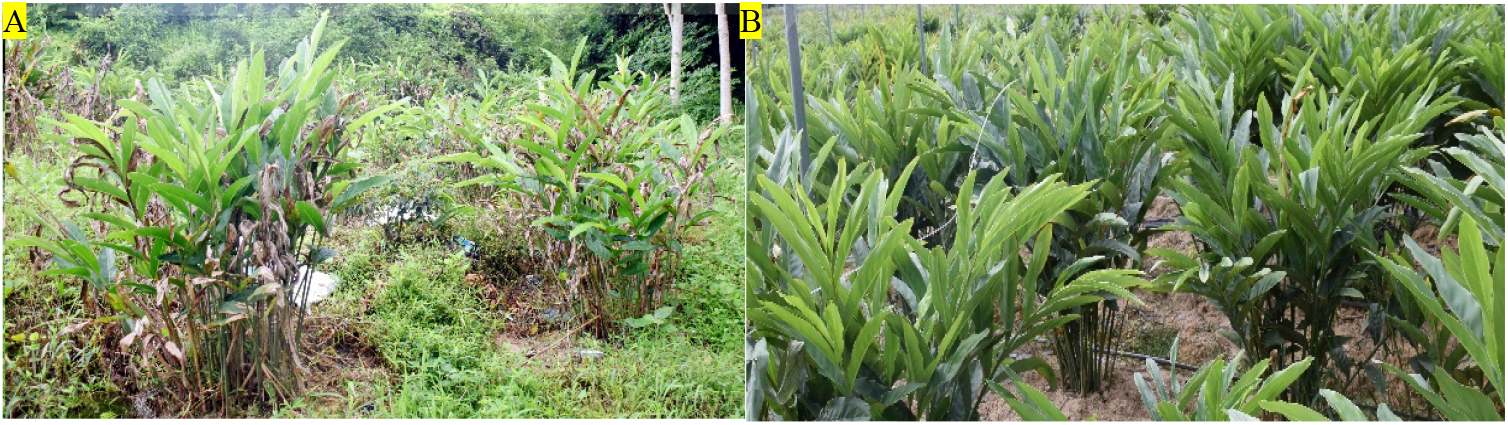
The growth of healthy and diseased plants of *A. galanga* in planting bases. A: HDK_B; B: HDK_J.

### Incidence Investigation

The surveyed planting plots of *A. galanga* were same to that of collected samples, and the investigation time was at during severe periods of bacterial wilt in May 2023.The incidence (%) and the severities of bacterial wilt were investigated, and the disease index (DI) was calculated. Incidence (%) = the numbers of infected plant clusters / the total numbers of investigated clusters ×100. The classification criteria for the severities of bacterial wilt of *A. galanga* refer to the national standards for the classification of tobacco bacterial wilt disease in China (GB/T 23224-2008). The severities of bacterial wilt are divided as follows: Grade 0: no diseases on the whole plant; Grade I: the scattered lesions at the bases of the stems, and narrow and discontinuous lesions, and no wilting of leaves; Grade III: the lesion fusion at the bases of the stems, and not more than 1/3 lesions around the stem circumference, and 1/3 lightly wilting leaves, and a few green withering in leaves of the lower parts of the plants; Grade V: more than 1/3 areas of merging lesions around the stem circumference, and 1/2 green withering leaves; Grade VII: all lesions around the stem circumference, and more than 2/3 green withering leaves; Grade IX: green withering and dead in all leaves. Disease index (%) =∑ (the numbers of diseased plant clusters at all levels × severity at all levels) / (total numbers of investigated plant clusters × highest severity) ×100.

### Material Handling

The samples were processed in three batches, and five healthy and diseased stems and leaves were randomly selected in each sample. The stems were cut into 5 cm lengths, and the leaves were cut in the 5×5 cm sizes. The order of sterilization is as follows: the stems and leaves were placed in beakers containing 75% ethanol for 3 min, and then rinsed three times in sterile water, followingly, the materials were soaked in 0.1% mercuric acid by adding Tween-20 for stems 1.5 min and for leaves 2 min. During the immersion period, the stems and leaves were stirred continuously so that they were in full contact with the sterilizing solution, and were removed and rinsed with sterile water for 1 min each time for 3 times, finally, the samples were blotted dry with sterile filter papers for later.

### Isolation, Culture, Purification and Preservation of Strains

A total of 60 tissue blocks were cut from healthy and diseased material under aseptic conditions by cutting the leaves into 1.0 × 1.0 cm size and stems into 0.5 cm size. Every three tissue blocks were placed on PDA, and the tissue blocks were rolled or laid on the surface of PDA for 2 minutes in advance to avoid microbial contamination as a control. Endophytic fungi were cultured in healthy tissue blocks with PDA medium.PDA and TTC (Guangdong Hankai Microbial Technology Co., Ltd, China) medium were used for the isolation and culture of endophytic fungi and pathogenic bacteria in diseased tissue blocks. The petri dishes were incubated at 25 °C under constant temperature and humidity and observed on the next day. When mycelium grew on the cut surface of the tissue block, it was picked out from the edge for isolation. Strain purification was performed by single picking of hypha at the edge of the colony in the early stage, and single spore isolation was used after spore production. The purified strains were numbered and inoculated into 9 cm test tubes and stored under refrigeration at 2∼4 °C for strain classification and identification.

### Strain Identification

#### Morphological identification

After the activation of the preserved strains, the morphological observation and preliminary screening were carried out with ZEISS (Axio Scope) optical microscope. The identification contents mainly included colony morphology, hypha growth characteristics, conidial stalk and conidiospore morphology, sporulation structure and mode of sporulation, etc. Strains that were difficult to spore-producing were cultured for sporulation, and then identified after sporulation according to the above methods.

#### Molecular identification

The initial screened strains were identified using the fungal ITS amplicons as molecular marker, and the DNA was extracted using the fungal genome DNA extraction kit (Beijing Soleberg Technology Co., Ltd, China). The primers used for PCR amplification were ITS1 (5’-TCCGTAGGTGAACCTGCGG-3’) and ITS4 (5’-TCCTCCGCTTATTGATATGC-3’) (Beijing Soleberg Technology Co., Ltd, China), and the DNA ITS1/ITS4 sequences were amplified. PCR amplification systems were (50 μL) : DNA template (100 ng.·L^-1^)1 μL, primers ITS1 and ITS4 2 μL of each, gold Mix (green) 45 μL. The PCR amplification procedure was: pre-denaturation at 98 °C for 2 min, denaturation at 98 °C for 10 s, annealing at 55 °C for 10 s, extension at 72 °C for 90 s, a total of 35 cycles, and finally extended at 72 °C for 1 min, and the reaction was terminated at 4 °C. According to the ratio of adding 1 μL DNA sample loading buffer to 5 μL DNA sample, the two were mixed for later. The mixture were directly set sampling with DL2000 DNA Marker, and the samples were spotted in 1% agar gel. The target bands were checked by UV analyzer 30 min after electrophoresis, and the amplified products were sequenced by the Guangzhou Company, Shenzhen BGI Genomics Institute, China.

The measured ITS sequences were identified using the fungal Database (UNITE Database), and the sequences with no or ambiguous matches in the identification process were then compared by BLAST in NCBI’s GenBank and were downloaded. Clustal X software (version 1.83) was used for multiple sequence comparisons, and MEGA software (version11.0) was used for the identification of relatedness and the construction of phylogenetic tree by Maximum-likelihood (ML) method.

#### Data Processing and Calculations

The analysis of diversity indicators included: (1) Colonization rate (CR) reflects the abundance of endophytic fungi in the host, and CR(%) = (the numbers of tissue blocks isolated from which endophytes were isolated / the numbers of total tissue blocks) ×100; (2) Relative separation frequency (SF) reflects the occurrence frequency of a microbial species in the endophytic fungal community in the same material, and SF (%) =(the numbers of isolated microbial species / the total numbers of all isolated microbial species)×100; (3) Relative abundance (RA) refers to the proportion of the numbers of individual species in the colony to the total numbers of species in the community; (4) Shannon-Wiener index (*H’*) is used to compare and to analyze the species richness level of endophytic fungal communities between healthy and diseased materials, and the calculation formula as follows: 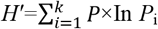, where *P*_*i*_ refers to a species percentage in all species, and higher values represent higher community diversity and complexity. (5) Simpson diversity index (*D*) was used to assess endophytic fungal community diversity, and the greater values are, the greater the community diversity is. The calculation formula as follows : *D*=1-Σ(*P*_i_)^2^, and where *P*_i_ represents the proportion of numbers of species in all species; (6) Margalef richness index (*R*) is calculated as follows: *R*=(S-1) / ln N, and where S represents the microbial numbers of species, and N represents the total numbers of samples; (7) Pielou evenness index (*J*) is calculated by the formula: *J* = *H′* / ln S, and where *H’* is Shannon-Wiener index, and *S* represents the numbers of species. Diversity indices among different samples were analyzed for significance of difference using one-way ANOVA test (*P* < 0.05), and the homogeneity of variance was tested by Levene’s test. When *P* < 0.05, the difference was significant. All statistical analyses were performed by Excel software and biostatistical software SPSS.

The difference analysis of community composition of endophytes includes: After molecular identification, the relative abundance of endophytic fungi species was analyzed at the genus and species level. Excel comma-separated value files (.csv) and ggplot2 software were used to draw the clustering heat maps and relative abundance stack maps of endophytic fungi species by R language.

## Results and Analysis

### Field Investigation of Bacterial Wilt

*A. galanga* rhizome is stout and well developed with a bright red surface, and the mother buds develop to form plant clusters ranging from 8 to 15 branches above ground per cluster. Taking plant cluster as the unit of investigation, the average incidence of bacterial wilt was 92.24% (Figure 2 A). The severities of aboveground branches varied from cluster to cluster, with older shoots having severe diseases and newer shoots having less or no. The highest disease index (DI) in aboveground branches of infected plant clusters was 87.11%, and average DI of all investigated plant clusters was 73.42%. The proportion of branches severities grade 0 is 6.32%, grade I 8.24%, grade III 17.55%, grade V 27.23%, grade VII 26.41%, and grade IX 14.25%. (Fig. 2 B, C).

**Figure 2.**
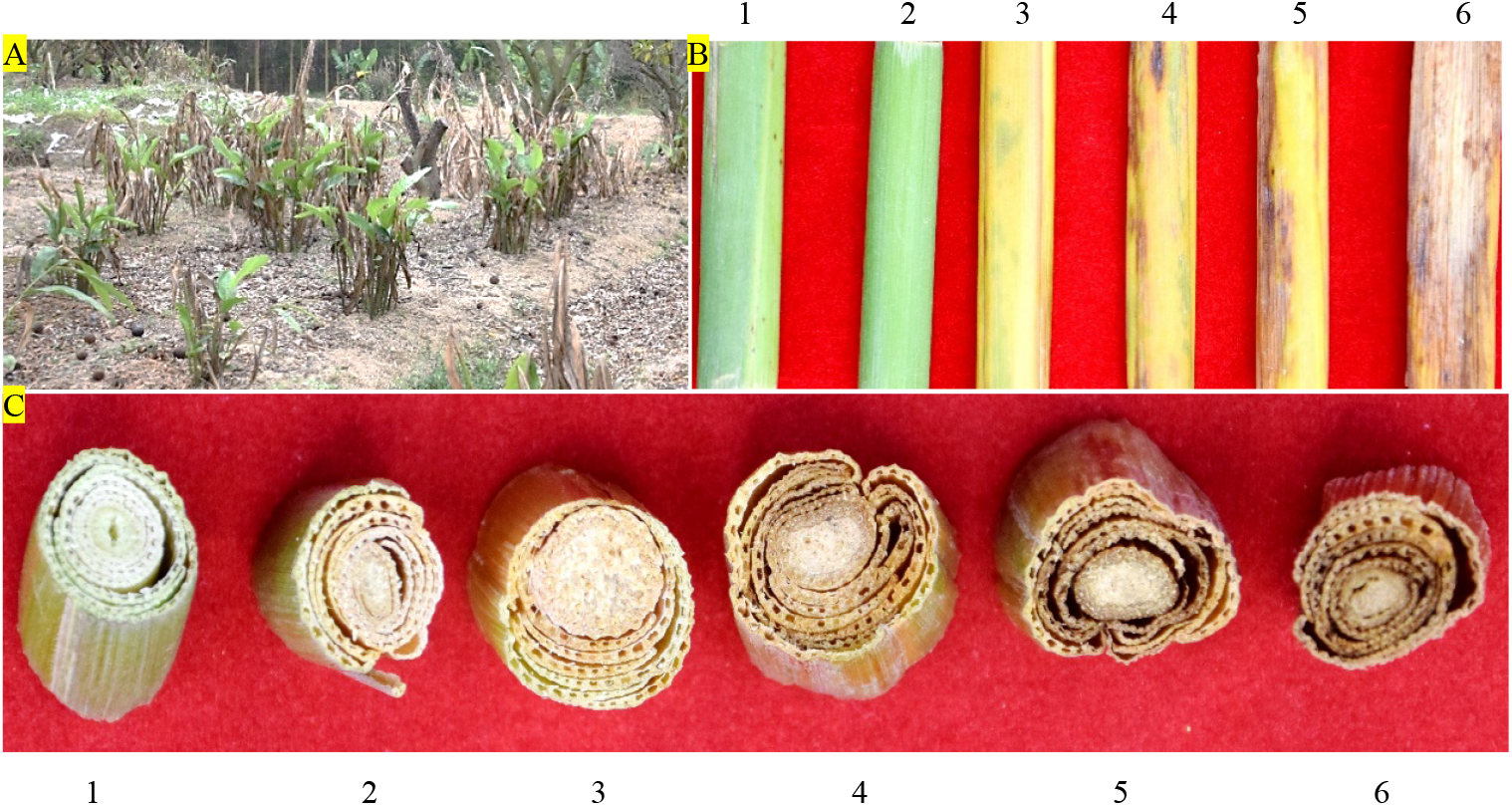
The incidence and severities of bacterial wilt in HDK_B. A: the incidence of bacterial wilt in HDK_B. B: the severities of bacterial wilt in the stems of HDK_B. C: the severities of bacterial wilt at basal parts of the stem transections in HDK_B. 1: HDK_J. 2: the severities at level I in HDK_B. 3: the severities at level III in HDK_B. 4: the severities at level V in HDK_B. 5: the severities at level VII in HDK_B. 6: the severities at level IX in HDK_B.

### Community Structure of Culturable Endophytic Fungi

Samples of healthy and diseased *A. galanga* were collected at different growth stages, and endophytic fungi were isolated and identified in 3 batches per stage. A total of 685 endophytic fungal strains were isolated, and a total of 556 strains were genetically characterized by fungal ITS genes after morphological characterization of colonies, hyphae, and conidia. Twenty-seven species of endophytic fungi were identified by ITS sequence alignment in the UNITE database and by BLAST software in GenBank database excluding duplicate species, of which 27 species were isolated from the healthy stems and leaves of *A. galanga*, and 8 species were isolated from diseased *A. galanga*. The 27 species are: *Alternaria* sp., *Epicoccum nigrum, E. andropogonis, E*. sp., *Pezicula* sp., *Colletotrichum boninense, C. gloeosporioides, C. fructicola, C*. sp., *Diaporthe subclavate, D*. sp., *Physalospora* sp., *Pestalotiopsis microspore, Nigrospora oryzae, N. musae, N*. sp., *Arthrinium arundinis, A*. sp., *Fusarium proliferatum, F. oxysporum, F*. sp., *Mycoleptodiscus indicus, M*. sp., *Xenoacremonium recifei, Trichoderma* sp., *Schizophyllum commune* and *Mucor fragilis*. Among them, the species of *Colletotrichum* were most abundant in healthy and diseased stems and leaves of *A. galanga*, and had the highest isolation frequency, indicating that the species of *Colletotrichum* are dominant endophytes. According to the latest mycological classification system, 27 species were categorized into 3 Phyla, 6 Classes, 9 Orders, 13 Families, and 16 Genera, of which, 16 were known species and 11 were unknown species (Table 1).

**Table 1.** The composition of community structure of endophytic fungi in *A. galanga* plant.

| Phylum | Class | Order | Family | Genus | Species |
| --- | --- | --- | --- | --- | --- |
| Ascomycota | Dothideomycetes | Pleosporales | Pleosporaceae | *Alternaria* | *A. sp.* |
|  |  |  |  | *Epicoccum* | *E. nigrum* |
|  |  |  |  |  | *E. andropogonis* |
|  |  |  |  |  | *E. sp.* |
|  | Leotiomyces | Helotiales | Dermateaceae | *Pezicula* | *P. sp.* |
|  | Sordariomycetes | Trichosphaeriales | Glomerellaceae | *Colletotrichum* | *C. boninense* |
|  |  |  |  |  | *C. gloeosporioides* |
|  |  |  |  |  | *C. fruticola* |
|  |  |  |  |  | *C. sp.* |
|  |  | Diaporthales | Diaporthaceae | *Diaporthe* | *D. subclavata* |
|  |  |  |  |  | *D. sp.* |
|  |  |  | Hyponectriaceae | *Physalospora* | *P. sp.* |
|  |  |  | Melanconidaceae | *Pestalotiopsis* | *P. microspora* |
|  |  |  |  | *Nigrospora* | *N. oryzae* |
|  |  |  |  |  | *N. musae* |
|  |  |  |  |  | *N. sp.* |
|  |  | Xylariales | Apiosporaceae | *Arthrimum* | *A. arundinis* |
|  |  |  |  |  | *A. sp.* |
|  |  | Hypocreales | Nectriaceae | *Fusarium* | *F. proliferatum* |
|  |  |  |  |  | *F. oxysporum* |
|  |  |  |  |  | *F. sp.* |
|  |  |  | Hypocreaceae | *Mycocleptodiscus* | *M. indicus* |
|  |  |  |  |  | *M. sp.* |
|  |  |  | Clavicipitaceae | *Xenoacremonium* | *X. recifei* |
|  | Xylonomycetes | Xylonomycetales | Xylonomycetaceae | *Trichoderma* | *T. sp.* |
| Basidiomycota | Agaricomycetes | Agaricales | Schizophyllaceae | *Schizophyllum* | *S. commune* |
| Zygomycota | Zygomycetes | Mucorales | Mucoraceae | *Mucor* | *M. fragilis* |

Further analysis of the taxonomic status of the 27 species showed that these species belonged to three phyla: Ascomycota, Basidiomycota and Zygomycota, with Ascomycota being the dominant phylum, accounting for 92.60%, Ascomycota for 3.70% and Zygomycota for 3.70%. At class level, 27 species belonged to 6 classes, including 4 classes of Ascomycota, namely Dothideomycetes, Leotiomycetes, Sordariomycetes and Xylonomycetes, and 1 class of Basidiomycota, Agaricomycetes, 1 class of Zygomycota, Zygomycetes. According to the included species numbers, the order of classes was Sordariomycetes (19 species) > Dothideomycetes (4 species) > Leotiomycetes, Xylomycetes, Agaricomycetes and Zygomycetes (1 species), of which Sordariomycetes was the dominant class, accounting for 70.37% of the total number of species. At the order level, 27 species belonged to 9 orders, including 7 orders of Ascomycota, namely Pleosporales, Helotiales, Trichosphaeriales, Diaporthales, Xylariales, Hypocreales and Xylonomycetales, Basidiomycota 1, Agaricales, Zygomycota 1, Mucorales. In order of the species included, the ranking order was Diaporthales (7 species) > Hypocreales (6 species) > Pleosporales and Trichosphaeriales (4 species) > Xylariales (2 species) > Helotiales, Xylonomycetales, Agaricales and Mucorales (1 species), of which Diaporthales was the dominant order, accounting for 25.93% of the total species. At the family level, 27 species belonged to 13 families, including 11 families of Ascomycota, namely Pleosporaceae, Dermateaceae, Glomerellaceae, Diaporthaceae, Hyponectriaceae, Melanconidaceae, Apiosporaceae, Nectriaceae, Hypocreaceae, Clavicipitaceae and Xylonomycetaceae, and 1 family of Basidiomycota, Schizophyllaceae, 1 family of Zygomycota, Mucoraceae. Among them, Pleosporaceae (4 species), Glomerellaceae (4 species) and Melanconidaceae (4 species) were the dominant families, accounting for 44.44% of the total species (Table 1).

### Analysis of Endophytic Fungal Diversity Index

The colonization and community composition of endophytic fungi were more related to the parts and health status of *A. galanga*. The colonization rate (CR) and diversity index (DI) of endophytic fungi in leaf tissues were higher than those in stems, and the CR and DI of endophytic fungi were higher in healthy *A. galanga* than those in diseased *A. galanga*, which were significantly different (*P*<0.05). As can be seen from Table 2: The leaf endophyte colonization rate was 130.94% of that of the stems, and the Shannon-Wiener index (*H′*), Simpson’s index (*D*) and Margalef’ s index (*R*) were HDK_J leaves (1.16, 0.87, 5.12) > HDK_J stems (0.82, 0.72, 4.22) > HDK_B leaves (0.57, 0.24, 2.97) > HDK_B stems (0.33, 0.14, 2.25). These results show that bacterial wilt disease had a great impact on the microenvironment of *A. galanga* stem and leaf tissues and destroyed endophytic fungal diversity and the stabilities of community structure, and comparatively, affecting the stem microenvironment was greater than the leaves. The relative separation frequency (SF) of dominant endophytic species under the same batch of tissue blocks from heathy and diseased *A. galanga* was that HDK_J (26.46%) was lower than HDK_B (50.69%), which was significantly different (*P*<0.05). The results indicated that the decrease of the relative abundance (RA) of endophytic fungal species in HDK_B caused the SF fluctuations of endophytic fungi in host tissues, and the SF was raised in HDK_B. At the same time, the Pielou index (*J*) of HDK_J leaves was higher than that of HDK_J stems and HDK_B stems and leaves, and the uniformity of HDK_J stems and HDK_B stems and leaves was not much different, indicating that the endophytic fungi species in HDK_J leaves were more evenly distributed (Table 2). The relative abundance clustering heat map of species showed that the community structure and composition of the endophytic fungi in the three groups of samples (HDK_J1, HDK_J2, and HDK_J3) from HDK_J were relatively stable, with little change in relative abundance of the 27 species, whereas the structure and composition in the three groups of samples (HDK_B1, HDK_B2, and HDK_B3) from HDK_B fluctuated greatly, with some of the stress-sensitive species disappearing, and some of the species turning out to be dominant endophytic fungi (Figure 3).

**Table 2.**
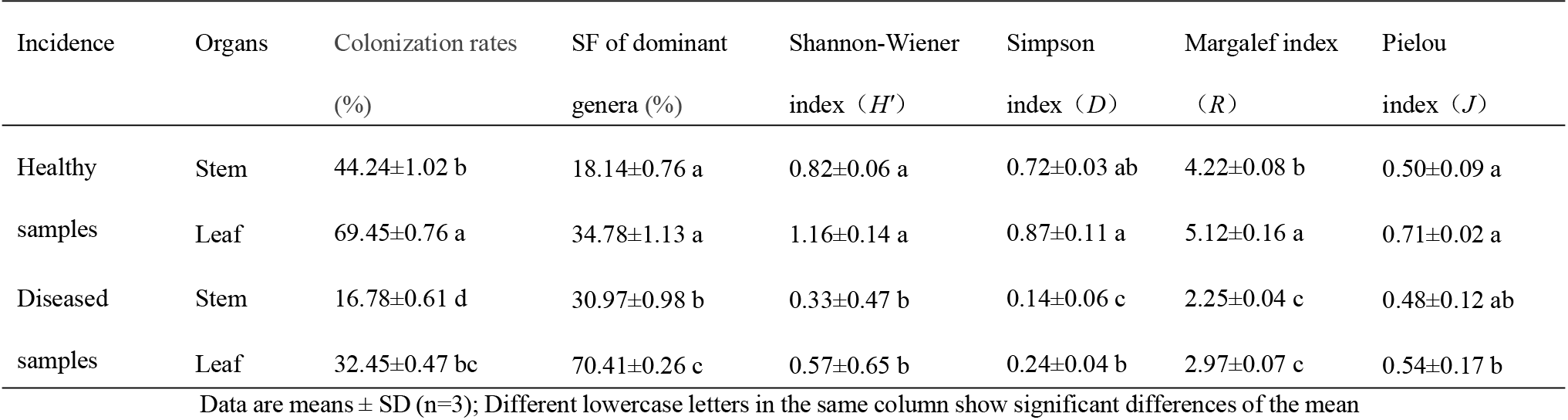
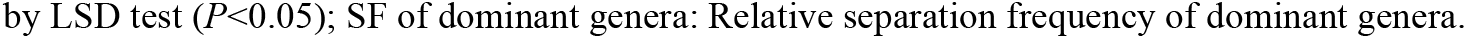
The diversity index of community structure of endophytic fungi in HDK_J and HDK_B.

| Incidence | Organs | Colonization rates | SF of dominant | Shannon-Wiener | Simpson | Margalef index | Pielou |
| --- | --- | --- | --- | --- | --- | --- | --- |
| | | (%) | genera (%) | index ($H'$) | index ($D$) | ($R$) | index ($J$) |
| Healthy | Stem | 44.24±1.02 b | 18.14±0.76 a | 0.82±0.06 a | 0.72±0.03 ab | 4.22±0.08 b | 0.50±0.09 a |
| samples | Leaf | 69.45±0.76 a | 34.78±1.13 a | 1.16±0.14 a | 0.87±0.11 a | 5.12±0.16 a | 0.71±0.02 a |
| Diseased | Stem | 16.78±0.61 d | 30.97±0.98 b | 0.33±0.47 b | 0.14±0.06 c | 2.25±0.04 c | 0.48±0.12 ab |
| samples | Leaf | 32.45±0.47 bc | 70.41±0.26 c | 0.57±0.65 b | 0.24±0.04 b | 2.97±0.07 c | 0.54±0.17 b |

Data are means ± SD (n=3); Different lowercase letters in the same column show significant differences of the mean
by LSD test ( $P<0.05$ ); SF of dominant genera: Relative separation frequency of dominant genera.

**Figure 3.**
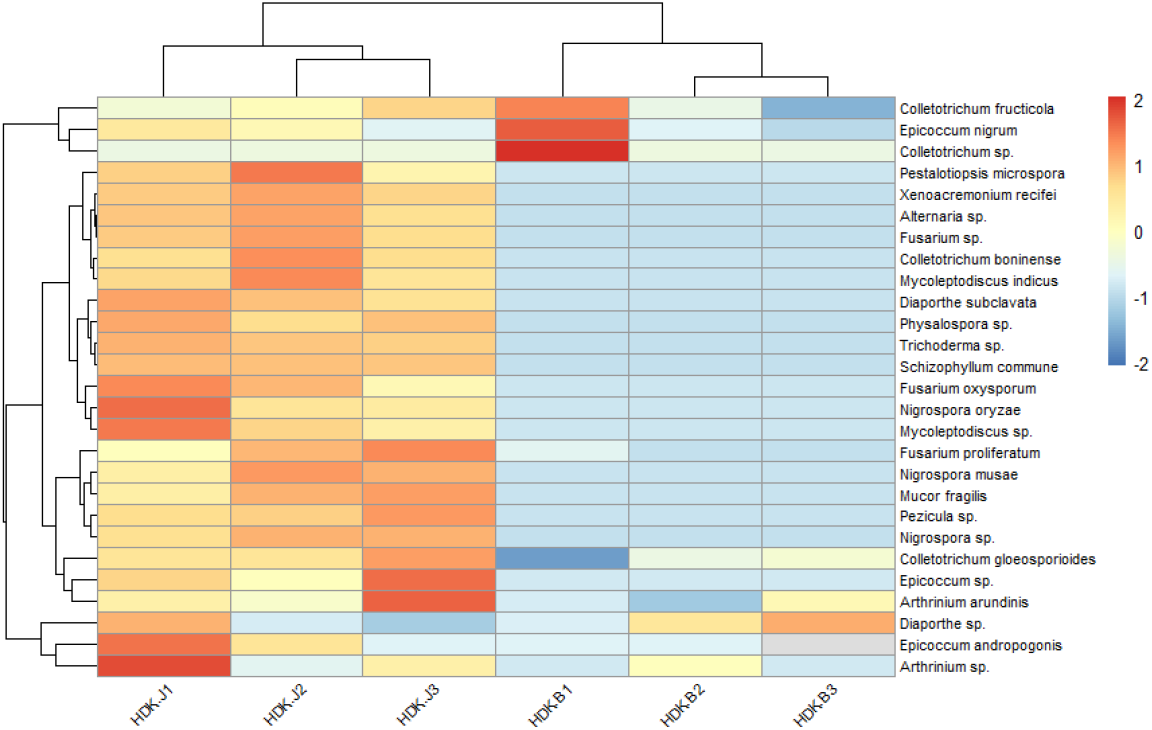
Abundance clustering heat map of endophytic fungi in HDK_J and HDK_B at species level. HDK_J1, HDK_J2 and HDK_J3: three samples of HDK_J. HDK_B1, HDK_B2 and HDK_B3: three samples of HDK_B.

### Differences of the Community Composition in Healthy and Diseased *A. galanga*

At the genus level, the endophytic fungi isolated from HDK_J belonged to fifteen genera, including thirteen genera in the Ascomycota, namely, *Alternaria, Epicoccum, Pezicula, Colletotrichum, Diaporthe, Physalospora, Pestalotiopsis, Nigrospora, Arthrinium, Fusarium, Mycoleptodiscus, Xenoacremonium* and *Trichoderma*, a genus in the Basidiomycota, *Schizophyllum*, and a genus in the Zygomycota, *Mucor*. The dominant genus was *Epicoccum* at the genus level with a relative abundance of 36.46%. Meanwhile, HDK_B was divided into 5 genera, namely, *Epicoccum, Colletotrichum, Diaporthe, Arthrinium*, and *Fusarium*, all of which were endophytic fungi of the Ascomycota. the dominant genus in HDK_B was *Colletotrichum*, with a relative abundance of 50.69%. Comparing the relative abundance of the HDK_J and HDK_B at the genus level, HDK_B endophytic fungi was only 38.93% of HDK_J. (Figure 4).

**Figure 4.**
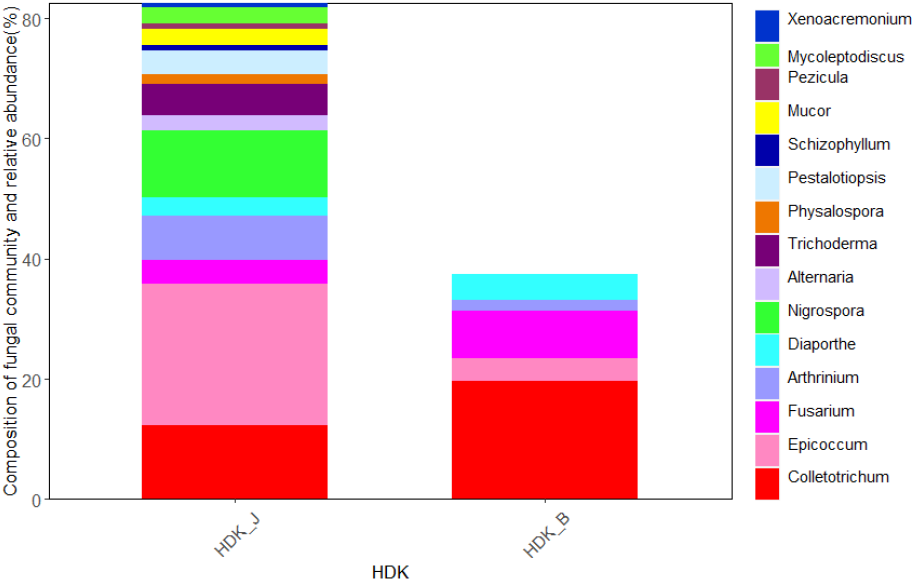
The community composition and relative abundance of endophytic fungi in HDK_J and HDK_B at genus level.

At the species level, the endophytic fungi species of HDK_J and HDK_B were significantly different. Among the 27 isolated endophytic fungi, HDK_J and HDK_B shared eight species of endophytic fungi, five known species, namely, *E. nigrum, C. gloeosporioides, C. fructicola, A. arundinis*, and *F. proliferatum* respectively, and three unknown species, namely, *Colletotrichum* sp., *Diaporthe* sp. and *Arthrinium* sp. These eight shared endophytic fungi had strong pigment-producing ability, among which the *E. nigrum* belonged to Leotiomycetes and the others belonged to Sordariomycetes. Most of the eight endophytic fungi were semi-living trophic fungi with low degree of specialization, and some of them were potentially pathogenic. The metabolism of the eight fungi in culture was vigorous, and the ability to produce secondary metabolites was strong, especially pigmented substances (Figure 5). On the other hand, the nineteen endophytic fungi isolated from HDK_J, including *Alternaria* sp., *E. andropogonis, Epicoccum* sp., *Pezicula* sp., *Colletotrichum boninense*, Diaporthe subclavata, *Physalospora* sp., *Pestalotiopsis microspora, Nigrospora oryzae, N. musae, N*. sp., *Fusarium oxysporum, F*. sp., *Mycoleptodiscus indicus, M*. sp., *Xenoacremonium recifei, Trichoderma* sp., *Schizophyllum commune* and *Mucor fragilis*, were sensitive to the stress of bacterial wilt, which affected the stability of endophytic fungal community structure in *A. galanga* (Figure 6).

**Figure 5.**
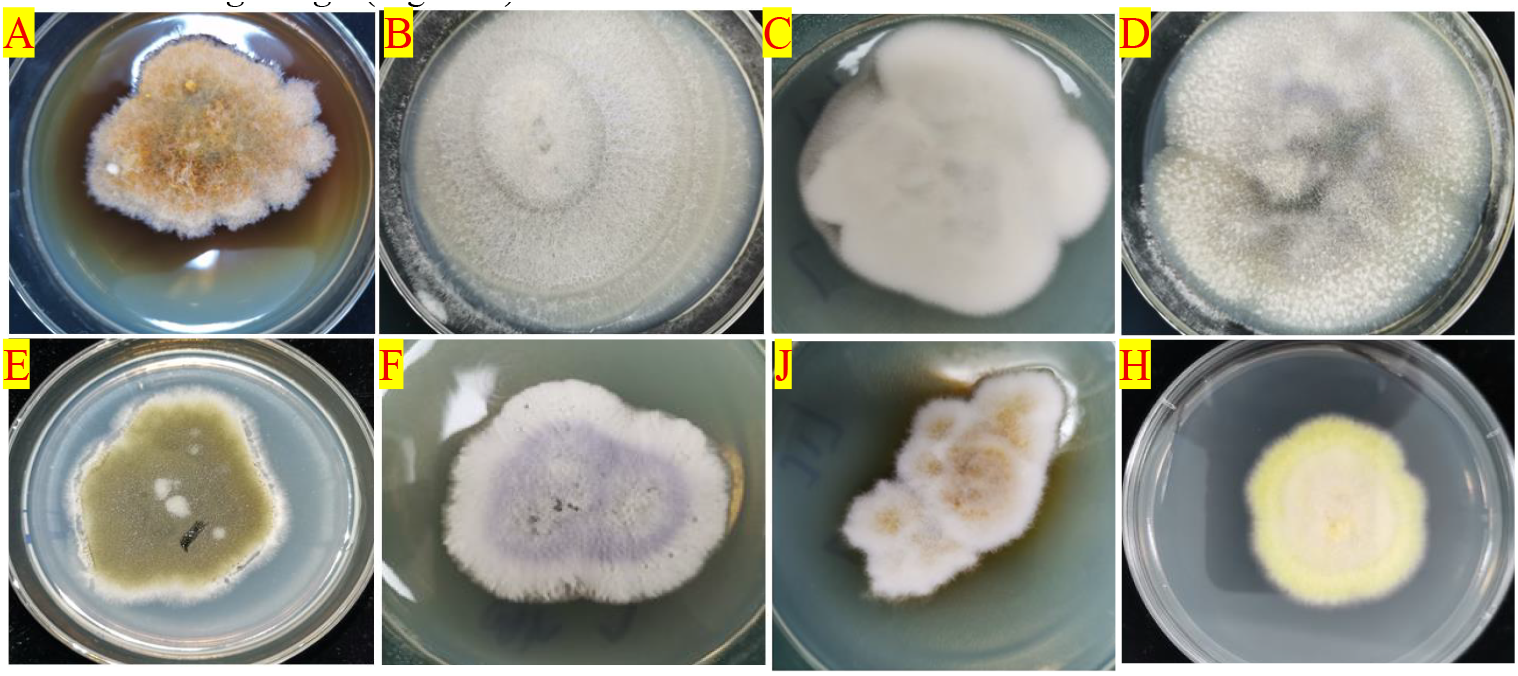
The colony morphology of the eight species of endophytic fungi producing pigments isolated from HDK_J/HDK_B cultured PDA medium. A: *Epicoccum nigrum*, B: *Colletotrichum gloeosporioides*, C: *C. fructicola*, D: *C*. sp., E: *Arthrinium arundinis*, F: *Fusarium proliferatum*, G: *A*. sp., H: *Diaporthe* sp.

**Figure 6.**
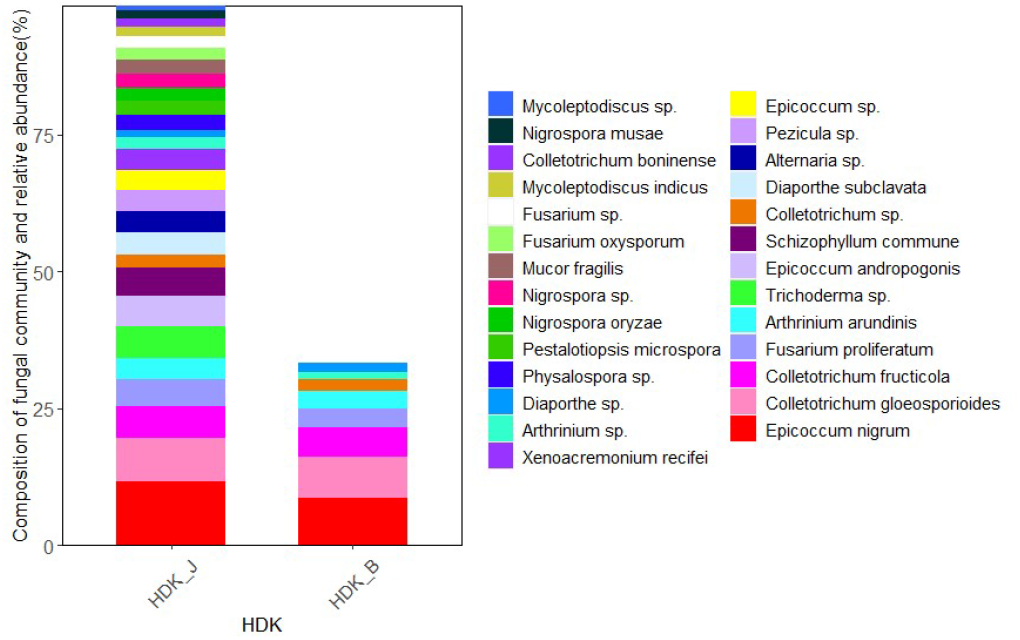
The community composition and relative abundance of endophytic fungi in HDK_J and HDK_B at species level.

### Difference of the Community Composition in the Stems and Leaves of *A. galanga*

The results of isolated endophytic fungi from the stems and leaves of *A. galanga* showed that the both organs had abundant endophytic fungi, but the relative abundance of leaves was higher than that of stems. Twenty-seven endophytic fungi were isolated from the stems and leaves of HDK_J, and the numbers of isolated species were the same, but the relative abundance of leaf organs was 124.23% higher than that of stem organs (Figure 7). Eight endophytic fungi were isolated from stems and leaves of HDK_B, with the same numbers of species in stems and leaves, but the relative abundance of leaf organs was 78.08% higher than that of stems (Figure 8).

**Figure 7.**
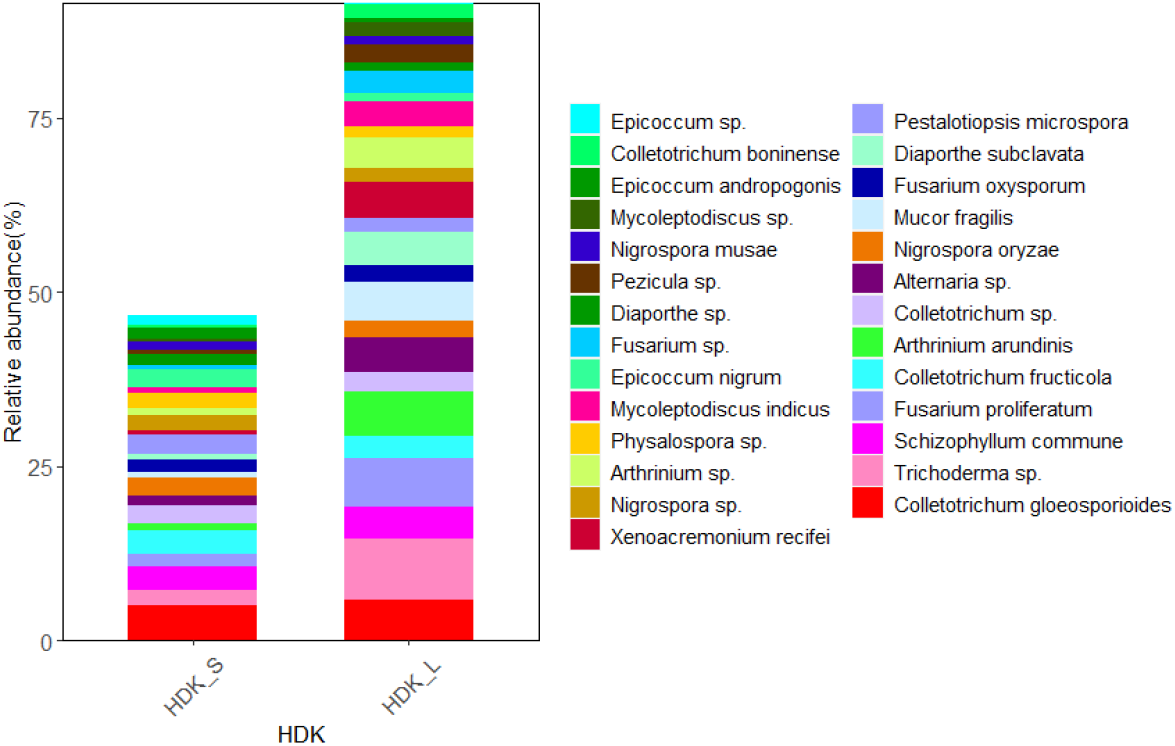
The composition of community and relative abundance of endophytic fungi in the healthy HDK_S and HDK_L.

**Figure 8.**
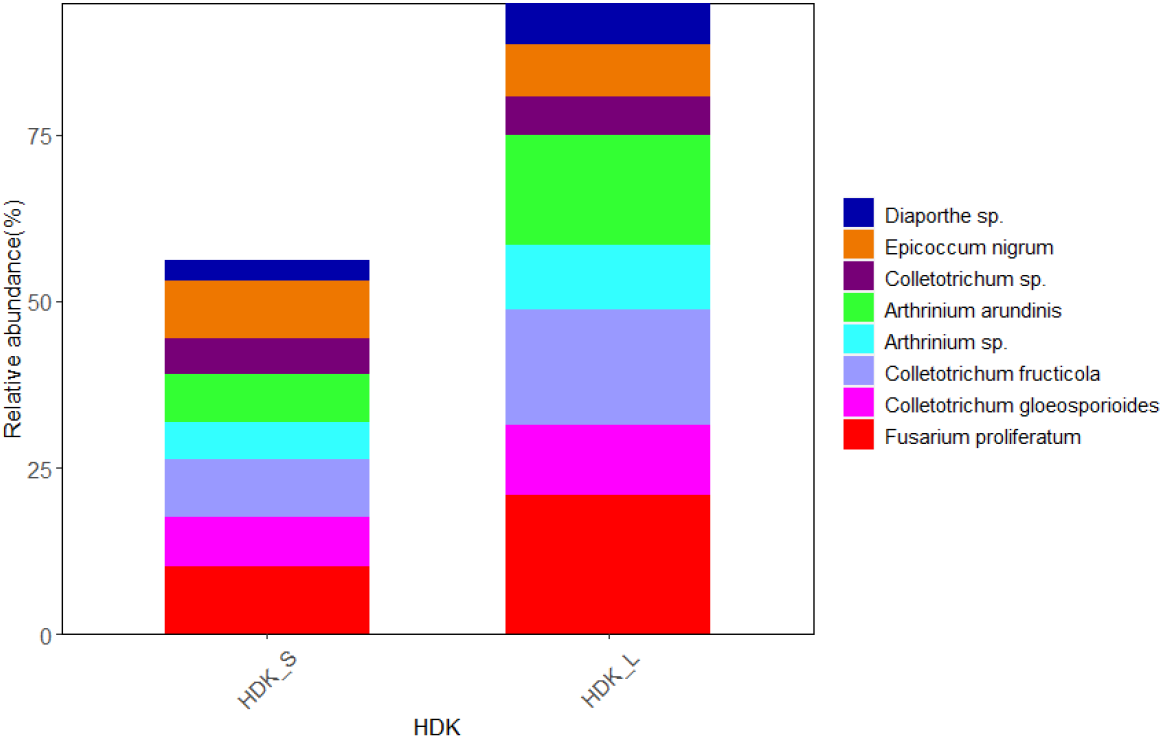
The composition of community and relative abundance of endophytic fungi in the diseased HDK_S and HDK_L.

### Phylogenetic Analysis of Endophytic Fungi of *A. galanga*

The determined gene sequences of species with ≥98% similarity to the sequences downloaded from the NCBI database were edited into fasta files using MEGA software, and the phylogenetic trees were constructed using the ML method (maximum likelihood method) with 1,000 bootstrap replications. The results showed that the phylogenetic trees of endophytic fungi in HDK_J could be divided into four branches, and the average genetic similarity among genera was 52.80%. Of these, *S. commune* and *M. fragilis* belonged to Basidiomycota and Zygomycota, respectively, and other species belonged to Ascomycota. *Mycoleptodiscus* was listed as a sub-branch alone in Ascomycota, the others were merged into a sub-branch. The longer lengths of branch of Ascomycota indicated a more complete and more variable genetic relationship between species. In addition, different genera, such as *Fusarium* and *Xenoacremonium, Alternaria* and *Epicoccum*, and *Arthrinium* and *Pestalotiopsis*, were closely related to each other, whereas *Fusarium* was relatively distantly related to *Arthrinium*. At the same time, *Physalospora* and *Trichoderma*, although being in the same branch, had fewer connections with each other, with a support rate of only 42% (Fig. 9). The endophytic tree of HDK_B had only one branch and was short. The average genetic similarity between genera was 84.59%. In addition to the close relation between species within *Colletotrichum, Colletotrichum* was relatively closely related to *Fusarium* and was relatively distantly related to *Diaporthe* and *Epicoccum* (Figure 10).

**Figure 9.**
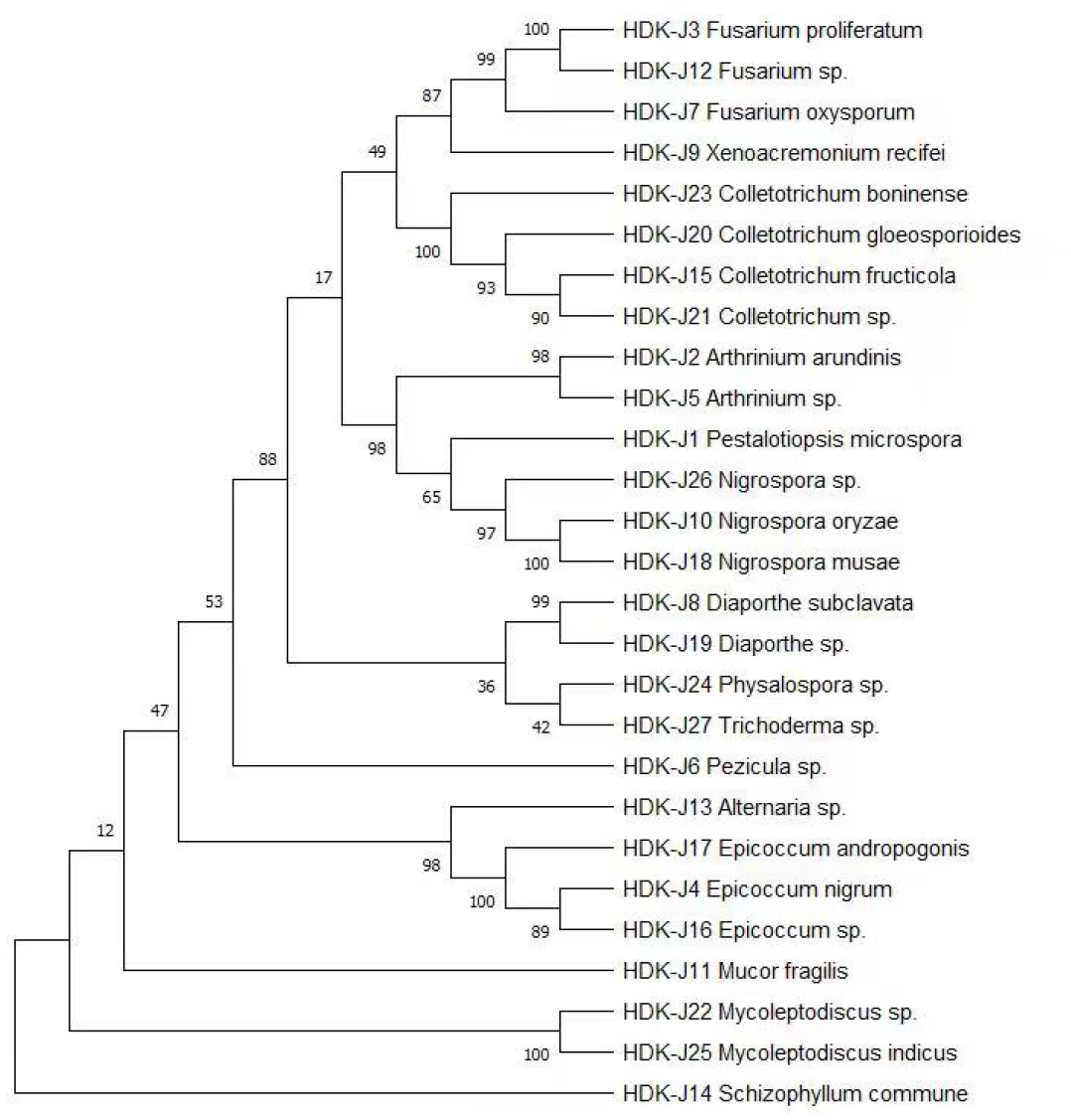
The phylogenetic relationships of endophytic fungi inferred from ML with MEGA in HDK_J.

**Figure 10.**
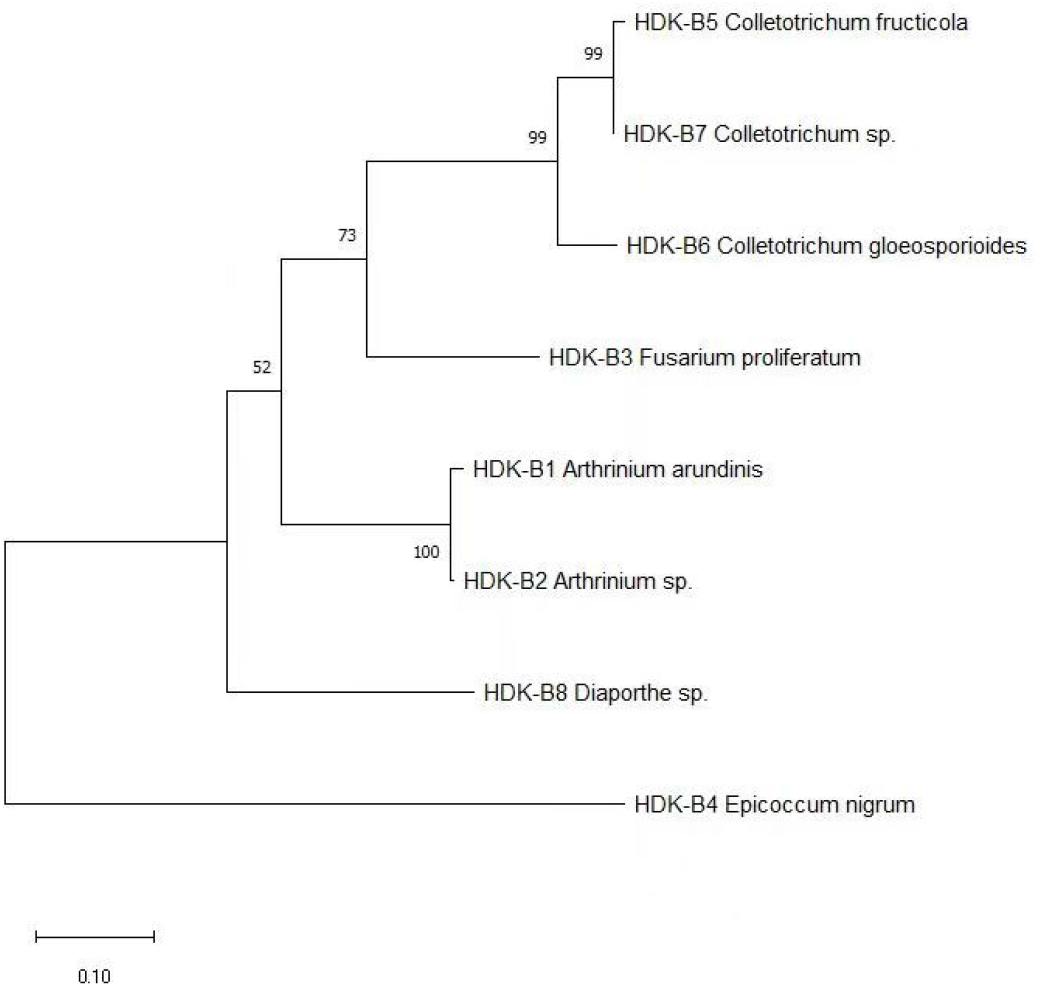
The phylogenetic relationships of endophytic fungi inferred from ML with MEGA in HDK_B.

## Discussion and Conclusion

Recruitment and community building of endophytes in host plants are multiply influenced by the external environment and internal physiological state [19,20]. External influences include biotic and abiotic factors, such as host diseases caused by bacterial infestation [21] and physiological pathologies caused by drought [22], waterlogging [23], soil salinization [24], atmospheric pollution [25], heavy metal overload [26] and nutritional deficiencies [27,28]. The effects of these factors on host endophyte communities are complex and variable. In general, these external factors drive the change of targeted endophyte recruitment and the spatial selection of endophyte colonization by influencing the host’s internal metabolic activities. The healthy hosts usually form a relatively stable endophyte community, through which assists the hosts to complete the normal growth and development as well as material accumulation, and resists various abiotic factor stress and exogenous pathogen infection in the form of a single microbial species or functional microbial groups in the long-term host-endophyte interactions. Targeted recruitment of endophytes by the host is closely linked to the soil, rhizosphere and leaf bulb microbial communities, and it has been shown that microbe-rich rhizosphere soils have higher diversity indices and endophyte abundance than microbe-deficient rhizosphere soils, which is particularly evident in perennial plants [29,30,31]. We found that it was difficult that endophytes were isolated from the tissue blocks of stems and leaves of Zingiberaceae plants that had been grown in pots by histocultured seedlings or rhizomes as asexual propagators, whereas samples from traditional planting areas with prolonged plantation had significantly higher diversity and abundance than those from non-traditional planting areas. Although the species and abundance of endophytes were closely associated with the microbial community attached to the body surface microenvironment of hosts, the community richness of endophytes was not consistent with that of other ecological niches, and the species and abundance of endophytes are change in different organs, with an overall decreasing trend of: soil microorganisms > rhizosphere microorganisms > phyllosphere microorganisms > endophytes, with particularly pronounced changes in bacterial community [32,33]. It has been shown that host physiological activities and metabolite accumulation are closely related to the structure composition of endophyte community, and that differences in host metabolites can lead to significant changes in the structure of the endophyte community, especially in metabolite-rich medicinal plants [34,35].

Similarly, the numbers and species of endophyte populations will be reduced in the hosts of disease due to the reduction in the accumulation of substances or the changes of chemical composition, which are caused by reduced metabolic activities or the changes of normal metabolic pathways of hosts that are infected by pathogens [36,37]. We found that the diversity and abundance of endophytic fungi in the stem and leaf tissues of diseased *A. galanga* were significantly lower than those of healthy *A. galanga*. At the species level, 27 endophytic fungi were isolated from healthy *A. galanga*, while 8 endophytic fungi were isolated from diseased *A. galanga*. Further analysis revealed that among 8 species of fungi isolated from both healthy and diseased hosts, 5 species of endophytic fungi, namely, *E. nigrum, C. gloeosporioides C. fructicola, A. arundinis* and *F. proliferatum*, belonged to hemibiotrophic fungi with a low degree of specialization, which are recruited to become endophytes in the non-host situation, and easily become pathogens after encountering the host, infecting the host and causing disease. Meanwhile, the 27 endophytic fungi were generally fast-growing and metabolically vigorous, and some species had strong pigment-producing ability. It has been demonstrated that the recruitment and colonization and functional microbiome assembly of species are not random, but selective, and are controlled by host transmembrane pattern recognition receptors (PRRs), which exhibit host-active selective recruitment patterns called host-associated molecular patterns (MAMPs), and that any factor affecting the expression of PRR transmembrane proteins causes changes in species abundance and diversity [38,39]. Recently, the PRR transmembrane proteins that interact with microorganisms have been found to be selective by knockout strategies, with different receptors recognising different species [40,41,42].

Plant endophytes combine with their hosts to form a holobiont, which assists the hosts in completing normal growth and development as well as in defense against external stresses. Thus, this holobiont, together with the microorganisms in the rhizosphere and the phloem layer, constitutes the plant microbiome, which characterizes the qualities of the hosts and maintains the health of hosts, and has the identity as “the second genome” of the hosts[43,44]. With the development of multi-omics analysis techniques including genomics, transcriptomics, proteomics and metabolomics, the function of the microbiome has begun to receive attention, and the molecular mechanisms of the interactions between the host and the microbiome, as well as between different microbiomes, have been explored from the perspective of microecology in order to provide a more in-depth and comprehensive understanding of the functions and compositions of micro-ecosystems and even of the ecosystem as a whole[45,46]. In general, different microbiomes of ecological niche do not interact with the host independently, but form a complex rhizosphere microbiome-endophyte microbiome-phyllosphere microbiome interaction network mediated by the hosts, or two or three synergistically affecting the hosts and exercising the roles of functional microbiomes. In the plant-soil-microbial cycling ecosystem, soil is an important factor affecting the recruitment, assembly and function of plant microbiome. Bever et al. [47] proposed the concept of “plant-soil feedback (PSF)”. According to this theory, there are complex checks and balances between plants and soil, between soil pathogens and rhizosphere microbiomes, and between soil pathogens and host endophytes. If the positive feedback effect continues, soil-borne diseases will be contained, and on the contrary, if the negative feedback effect continues, the index of microbial diversity will decrease, leading to the reduction of community function, which has been also confirmed by our experimental results. We constructed the phylogenetic trees of endophytes of healthy and diseased *A. galanga* using the ML method from the perspective of culturable endophytes, and found that the phylogeny of endophytes of healthy stems and leaves of *A. galanga* had four major branches, belonging to Ascomycota, Basidiomycota and Zygomycota respectively, which were Significant differences, and there were many longer sub-branches under the Ascomycota, suggesting that the genetic relationship on the branch of Ascomycota was more complete and the richer diversity. In contrast, there was only a branche of Ascomycota in the phylogenetic trees of the endophytic fungi in diseased stems and leaves of *A. galanga*, and there were also few sub-branches under Ascomycota, with close genetic relationships and small differences among genera, and a significant decrease in the diversity index, which was basically consistent with the study results of the sclerotinia rot disease (*Sclerotinia sclerotiorum*) by Zhang et al. [48]. Moreover, the host endophytic microbial community system includes not only endophytic fungal community, but also endophytic bacterial community and endophytic actinobacterial community, and it would be a meaningful work to explore how the synergistic effect among different community affects host development by using multi-omics techniques, especially in the field of medicinal plants. With the discovery of more and more new active endophytic strains through the genome sequencing, targeted construction of functional recombinant endophytic communities and artificially tailored synthetic microbial communities contributes to or promotes host growth and development and component accumulation.

